## Supplemental Figures for "Chemotherapy activates inflammasomes to cause inflammation-associated bone loss"

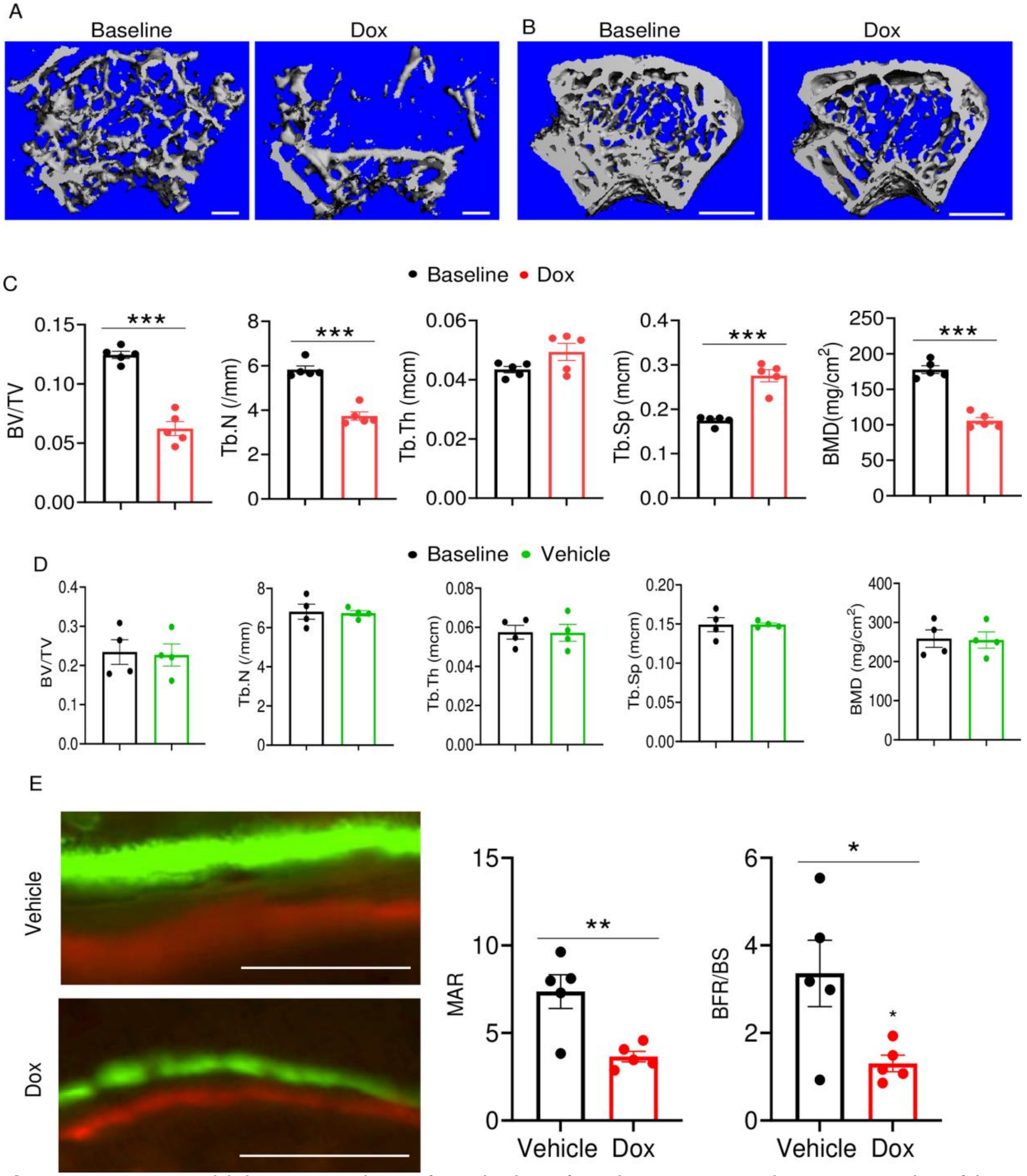


**Figure S1**: Doxorubicin causes bone loss in female mice. Femurs from 10 weeks old WT female (A, C) and male (B, D) mice were analyzed by VivaCT before (baseline) and 4 weeks after a single i.p. injection of 5 mg/kg doxorubicin (A-C) or vehicle (D). For dynamic histomorphometry, male mice were i.p. injected with calcein green and 6 days later with alizarin red, and were euthanized 2 days after the second injection. The tibias were used for the analysis. Cross sections of 3D reconstructions of trabecular (A; scale bar: 200 μm) and trabecular and cortical (B; scale bar: 500 μm) bones. (C, D) Bone parameters. (E) Dynamic bone parameters. Scale bar: 50 μm. Data are mean ± SEM. N=4-5 mice/group. *$P < 0.05$; **$P < 0.01$; ***$P < 0.001$. BFR/BS, bone formation rate/bone surface; BMD bone mineral density BV/TV, bone volume/total volume; MAR, mineral apposition rate; N.Oc/BS, OC number/bone surface; ns, not significant; OC, osteoclast; Tb.N, trabecular number; Tb.Sp, trabecular separation; Tb.Th, trabecular thickness.

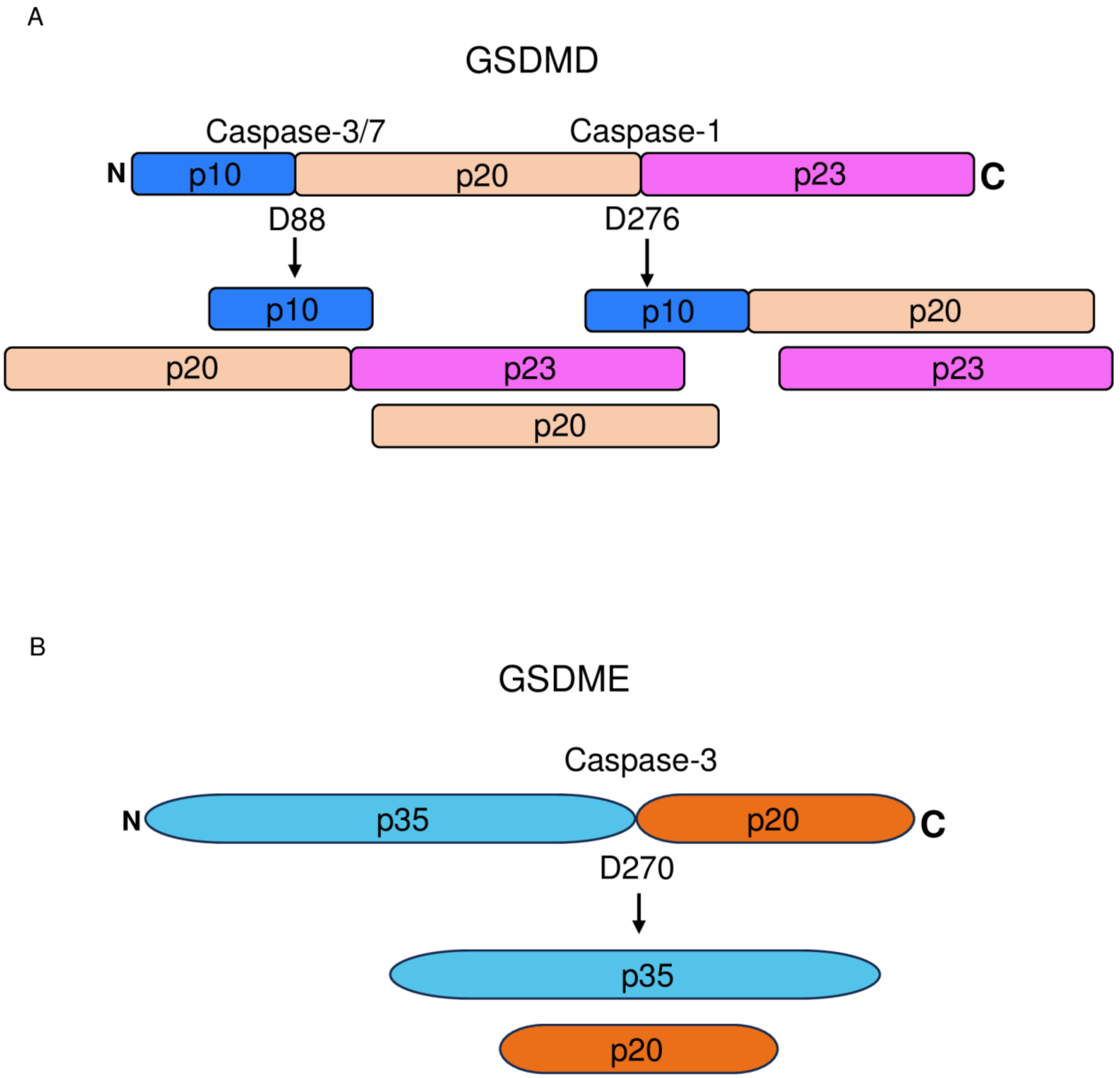


**Figure S2**: (A) Cleavage sites of murine GSDMD by caspase-1 and caspase-3. (B) Cleavage sites of murine GSDME by caspase-3.

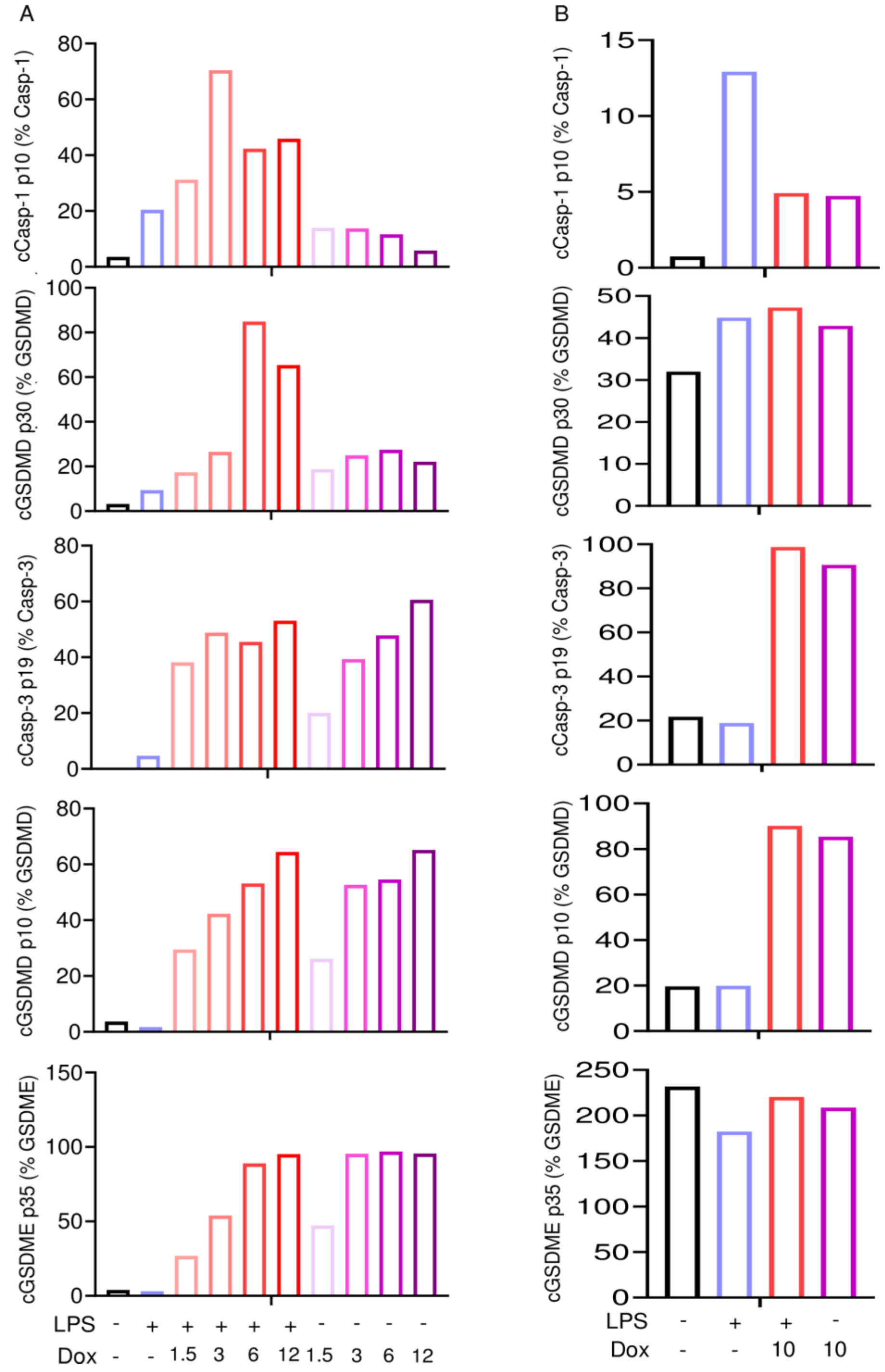


**Figure S3**: **Doxorubicin activates inflammasome –dependent and –independent pathways in macrophages and neutrophils.** WT BMMs and neutrophils were left untreated or primed with LPS for 3 hours, then treated with various doxorubicin concentrations for 16 hours. Whole cell lysates were analyzed by immunoblotting. (A) Quantitative data from Fig. 4. (B) Quantitative data from Fig. 5.

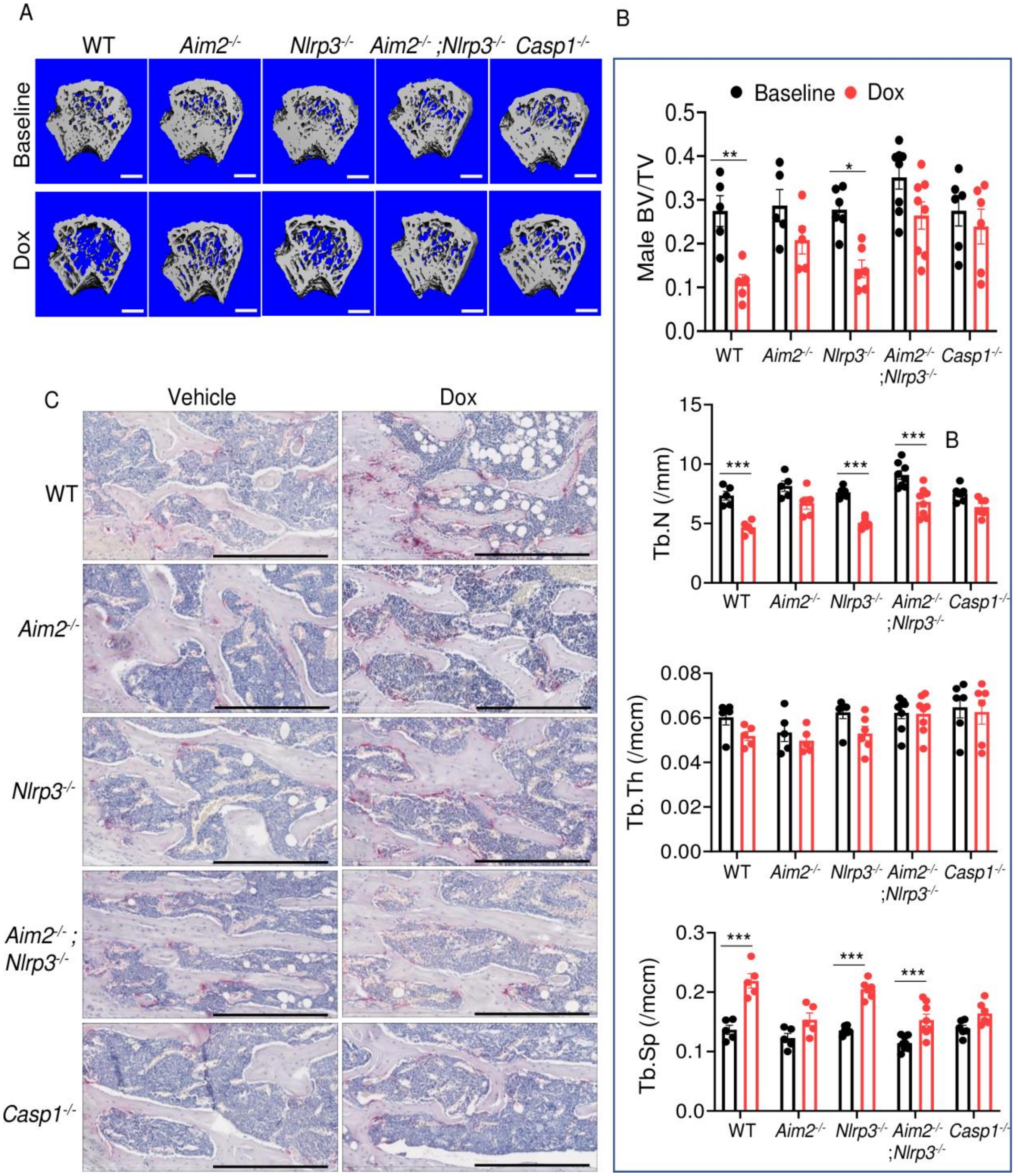


**Figure S4**: AIM2 and NLRP3 inflammasomes are involved in bone-damaging effects of doxorubicin in male mice. (A, B) Femurs from WT, *Aim2*$^{-/-}$, *Nlrp3*$^{-/-}$, *Aim2*$^{-/-}$;*Nlrp3*$^{-/-}$ or *Casp1*$^{-/-}$ male mice were analyzed by VivaCT before (baseline) and 4 weeks after a single intraperitoneal injection of 5 mg/kg doxorubicin. Cross sections of 3D reconstructions of trabecular and cortical (A; scale bar: 500 µm) bones. (C) Bones from WT, *Aim2*$^{-/-}$, *Nlrp3*$^{-/-}$, *Aim2*$^{-/-}$;*Nlrp3*$^{-/-}$ or *Casp1*$^{-/-}$ male mice were analyzed 4 weeks after a single intraperitoneal injection of vehicle or doxorubicin; specimens were stained for TRAP activity. Scale bar: 400 µm. Data are mean ± SEM. N=5-8 mice/group. ***$P < 0.001$. Two-way ANOVA was used. BV/TV, bone volume/total volume; BMD bone mineral density; Tb.N, trabecular number; Tb.Th, trabecular thickness; Tb.Sp, trabecular separation; WT, wild-type.

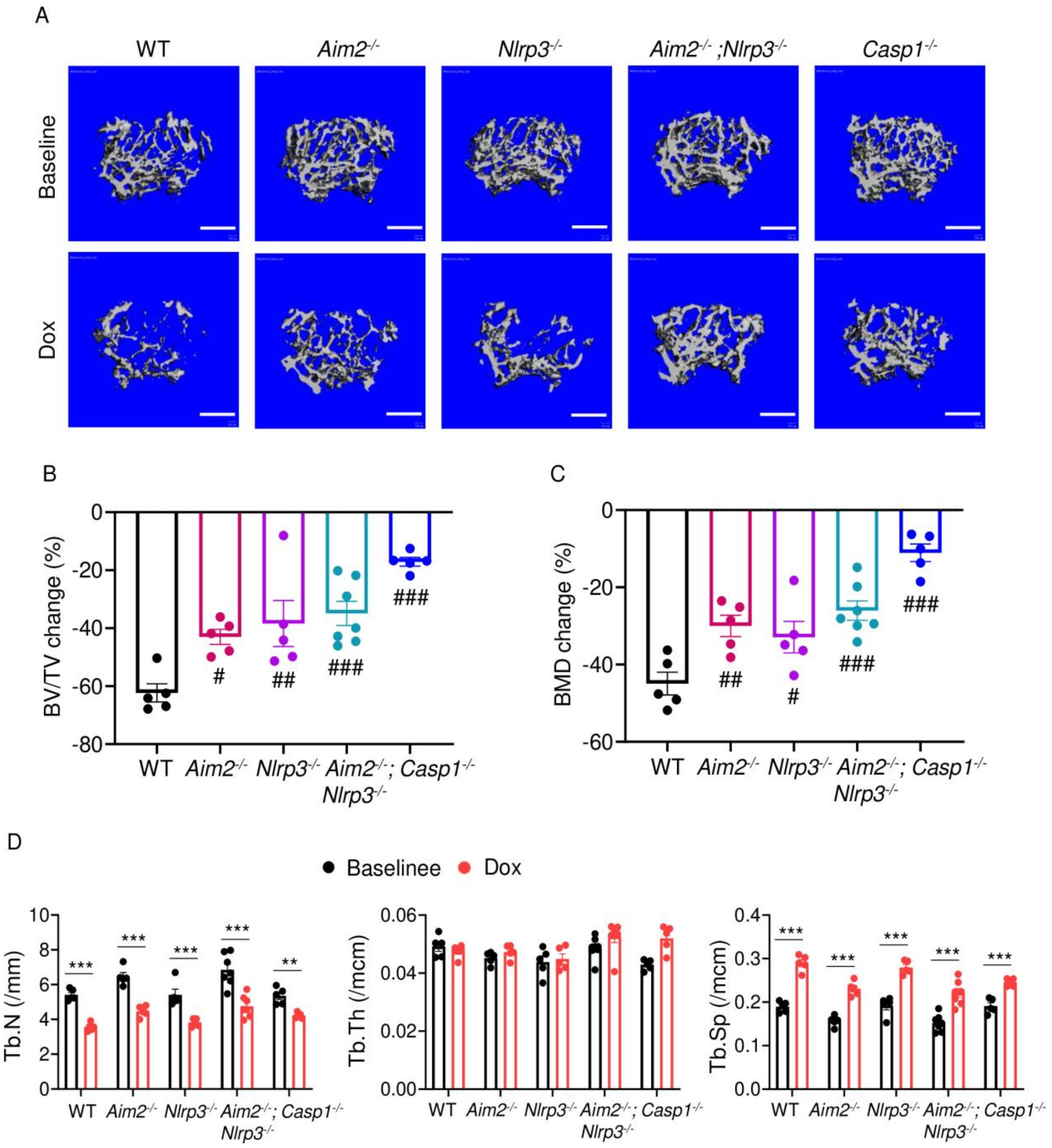


**Figure S5**: AIM2 and NLRP3 inflammasomes are involved in bone-damaging effects of doxorubicin in female mice. Femurs from WT, *Aim2*$^{-/-}$, *Nlrp3*$^{-/-}$, *Aim2*$^{-/-}$;*Nlrp3*$^{-/-}$ or *Casp1*$^{-/-}$ female mice were analyzed by VivaCT before (baseline) and 4 weeks after a single intraperitoneal injection of 5 mg/kg doxorubicin. (A) Cross sections of 3D reconstructions. (B) BV/TV changes. (C) BMD changes. (D) Additional bone parameters. Data are mean ± SEM. N=5 mice/group. **$P < 0.01$; ***$P < 0.001$; $^{\#}P < 0.05$; $^{\#\#}P < 0.01$; $^{\#\#\#}P < 0.001$. One-way ANOVA (B,C) and Two-way ANOVA (D) were used. Scale bar: 500 μm. BV/TV, bone volume/total volume; BMD bone mineral density; Tb.N, trabecular number; Tb.Th, trabecular thickess; Tb.Sp, trabecular separation; WT, wild-type.

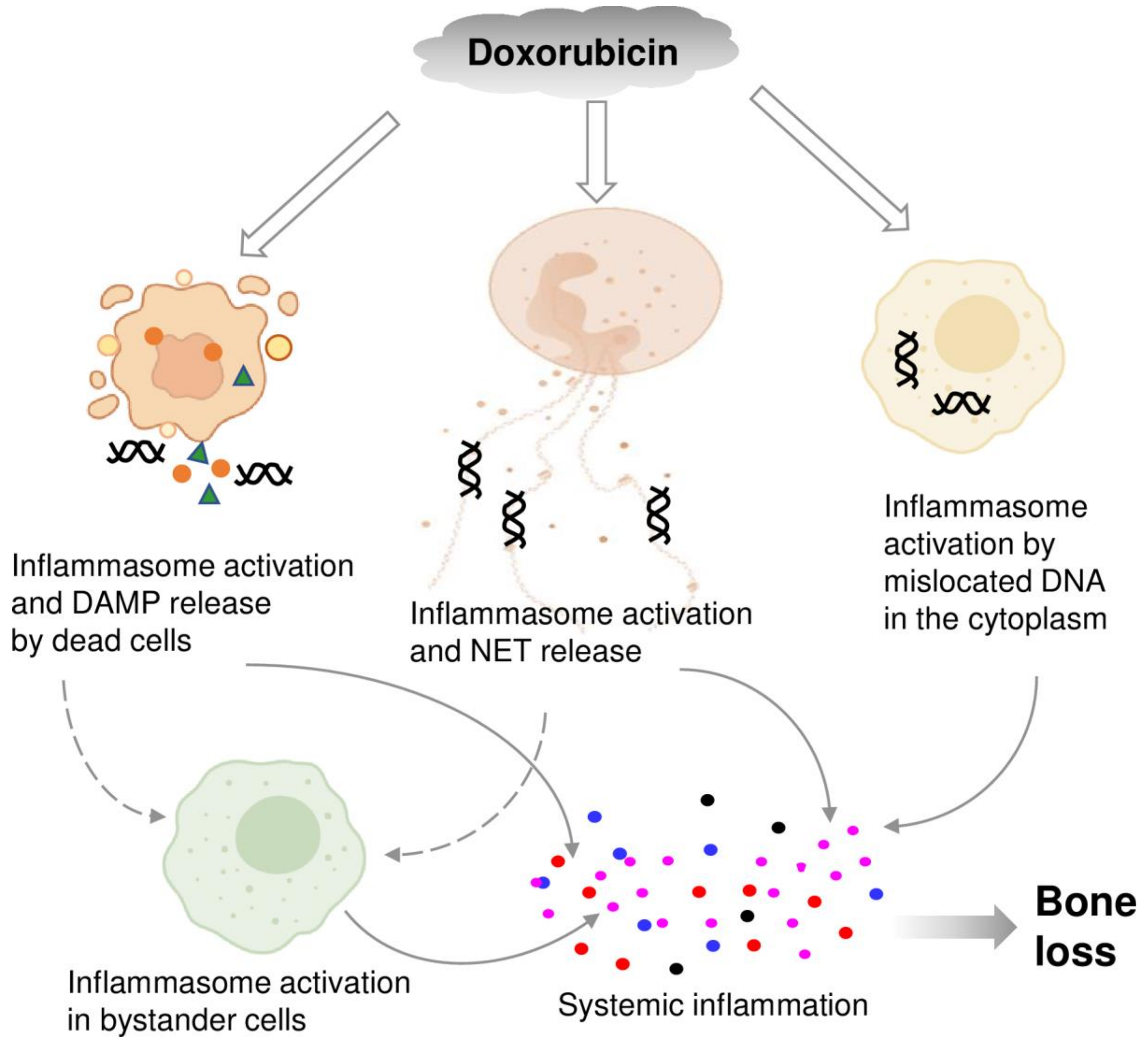


Fig. S6:

- Double line arrows: direct effects of dox on target cells
- Solid line arrows: direct contribution to systemic inflammation
- Broken line arrows: indirect contribution to systemic inflammation

(created with Biorender.com)
